# A long-hidden prevalent PCR artifact and its application in DNA library screening

**DOI:** 10.64898/2026.07.17.737729

**Authors:** Xinglin Jiang, Tue Sparholt Jørgensen, Fan Wang, Annette Lien, Tetiana Gren, Lachlan Jake Munro, Haibo Zhang, Tilmann Weber

## Abstract

PCR artifacts can lead to aberrant DNA products and confound result interpretation, posing significant challenges in research and diagnostics. Traditionally, it has been blamed to misannealing of primers on unintended sites. However, by sequencing, we find that most PCR byproducts and artifacts actually have both primers bound correctly but lack a middle part of the target, a phenomenon we named as PCR leaping. The leaping occurs at random positions, is independent of sequence repeats and requires a single piece template, indicating an unprecedented mechanism. It is observed with various polymerases, template qualities, GC contents, and PCR programs, thus necessitating a reset of our PCR understanding and careful consideration in future studies, for example in CRISPR editing validations. On the other hand, it provides a simple way of DNA end pairing, based on which we develop a high throughput DNA library screening method for natural product discovery at substantially reduced cost.

## Introduction

Polymerase chain reaction (PCR) employs a DNA polymerase and a pair of primers to selectively amplify a target DNA sequence ^1^. Since its invention in 1984, it has become a widely used and often indispensable technique in many fields ^2,3^. A fundamental requirement in PCR is that the product faithfully represents the original DNA target. Nonspecific amplification can lead to false judgments in PCR-based diagnostics, introduce artifacts in DNA sequencing, and aberrant products in cloning and genetic engineering ^4^. Traditionally, it has been attributed primarily to mispriming, where primers bind to off-target template sites ^5-11^. Once extended, shorter by-products amplify more efficiently than the intended amplicon and can rapidly dominate the exponential amplification ^4^. Based on this model, people have developed many strategies to improve PCR including primer sequence optimization^4^, primer modification^9,12-16^, hot-start PCR^7^, touchdown PCR^8^, nested PCR^17^, and PCR additives^18^,^6^. Hot-start and touchdown PCR are routinely used, as they are effective and easy to perform.

In this study, our sequencing results revealed that most nonspecific amplification products are rather generated from a previously unrecognized phenomenon that we named as PCR leaping. Unlike previously described chimeric PCR artifacts^25^, PCR leaping occurs independently of sequence repeats and therefore cannot be prevented simply by avoiding repetitive sequences in the template. Structurally, the leaping product is indistinguishable from a genuine long-deletion and therefore can be particularly problematic for PCR-based sequence validation, for example in CRISPR genome editing studies.

Meanwhile, it offers an alternative to the traditional DNA end-pairing methods that rely on circulation, random fragmentation, and ligation^19,20^. Combining it with primer barcoding and NGS sequencing, we developed a high throughput DNA library screening platform for isolating biosynthetic gene clusters (BGCs) from BAC libraries that has contributed to the discovery or study of several new natural products. This method may help revive the antibiotic discovery pipeline by facilitating access to cryptic BGCs and avoiding compound rediscovery.

## Results

### Most artifact products have both primers bound correctly but a middle part of the target leaped over

PCR is a fundamental technology and has been extensively optimized. However, a systematic evaluation of PCR results using modern sequencing has not been reported. To assess the prevalence and potential impact of PCR artifacts on current and emerging PCR-based technologies, we systematically analyzed PCR products by deep sequencing. First, we checked the byproducts in regular PCR reactions. Circular pUC19 plasmid and linear lambda DNA were amplified with ten primer pairs each using recommended annealing temperature and extension time. Full-length products were all successfully amplified (Figure 1a), appearing as bright single bands. Faint smears in the low molecular weight region were recovered from the gel and examined by illumina sequencing. Classic mispriming products were detected but surprisingly a new type of byproduct appeared six times more frequently (Figure 1b). This artifact, which we named as PCR leaping, is composed of the two target terminal sequences, suggesting that both primers have annealed and extended correctly. However, the middle part of the target is missing, leaving a junction point formed between the two leaping ends from the template (Figure 1c and d).

**Figure 1.**
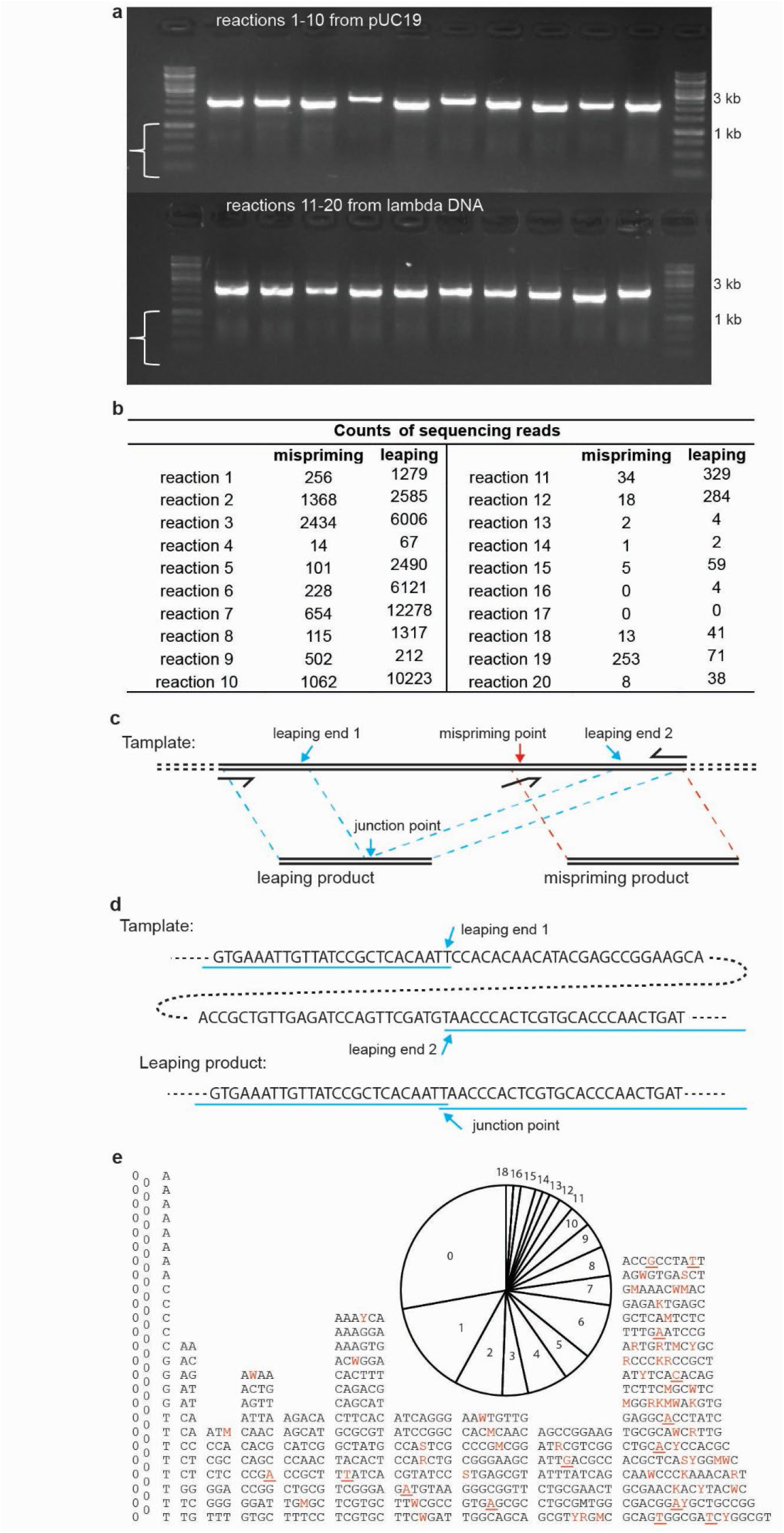
Analysis of byproducts in regular PCRs. **a**, Products of independent reactions were resolved on agarose gel. The expected amplicons range from 1.9 to 2.3 kb. The nonspecific byproducts (indicated by brackets) were excised and examined by Illumina sequencing. **b**, Read counts of different byproducts observed in each reaction. **c**, Schematic of mispriming (right) and PCR leaping (left) products. **d**, An example of a PCR leaping product. A one bp overlap was found at the junction point. **e**, Junctions from all dereplicated reads. A value of 0 indicates no overlap. Mismatched bases within the overlaps are shown using IUB codes in red; deletions are indicated by red underlined letters. The pie chart shows the distribution of overlap lengths.

Junctions with no overlap between the two leaping ends were most prevalent, followed by those with 1 bp overlaps (Figure 1e). They accounted for approximately half of the cases. It is still less than the theoretical value if leaping occurs at completely random positions, suggesting microhomology is not a prerequisite for the leaping but may facilitate it. Moreover, by reducing extension time and annealing temperature, nonspecific products became the dominant bands on the gel and most of them were confirmed to be leaping products by sequencing (supplementary Figure 1).

Next, we examined long-range PCR reactions where full-length targets are challenging to amplify, allowing artifact bands to dominate. BAC plasmid DNA as template was amplified at different annealing temperatures by a pair of universal primers flanking the cloning site. In addition, forced mispriming was performed for comparison, in which one of the specific primers was omitted, as in single primer PCR^21^. At the recommended annealing temperature of 60°C or a lower temperature of 50°C, two specific primers generated artifact bands which were determined to be leaping products by sanger sequencing. In contrast, no clear band was generated when only one specific primer was used. At a further reduced temperature of 40°C, bands appeared in the one specific primer reactions and were confirmed to be mispriming products. (supplementary Figure 2), suggesting mispriming requires lower annealing temperature than PCR leaping.

More PCR conditions were tested, including different templates (purified plasmid DNA, crude extracts, *E*.*coli* cells directly from agar plates, or PCR products), GC contents ranging from 45% to 72%, various polymerases (non-proofreading Taq, proofreading Q5, or OneTaq, a blend of Taq and Deep Vent®), primer with or without hairpin protection, and PCR programs with or without hot-start and touchdown. In all cases, most of the unspecific bands were found to be leaping products by sanger sequencing. (supplementary figure 2 to 7). The exonuclease activity of proofreading polymerase can repair 3’ end mismatch and reinitiate extension ^23,24^. Here we show that it is neither required nor prohibitive for the PCR leaping.

### PCR Leaping is a result of abnormal extension within the same template

When PCR is performed on a mixture of homologous templates, chimeric products can be generated through an artifact known as “template switching” or “jumping PCR”^25^. This occurs when the 3’ end of a prematurely terminated extension product anneals to a different template with sufficient sequence homology in a subsequent PCR cycle and is then extended by DNA polymerase ^26-28^. To check if PCR leaping is a result of “jumping PCR” between two sections of the same template, we compared reactions from a single contiguous template versus two separated fragments (figure 2a). Different template concentrations from 5E-7 ng/ μl up to 0.005 ng/ μl were tested. Short extension time was used to enrich nonspecific amplifications. As shown in the gel photo (supplementary figure 8), two-piece template required higher concentration than single piece template to generate nonspecific bands. Sanger Sequencing showed that PCR leaping products were only generated from the single piece template, and double-piece template generated only primer mispriming products (figure 2b).

**Figure 2.**
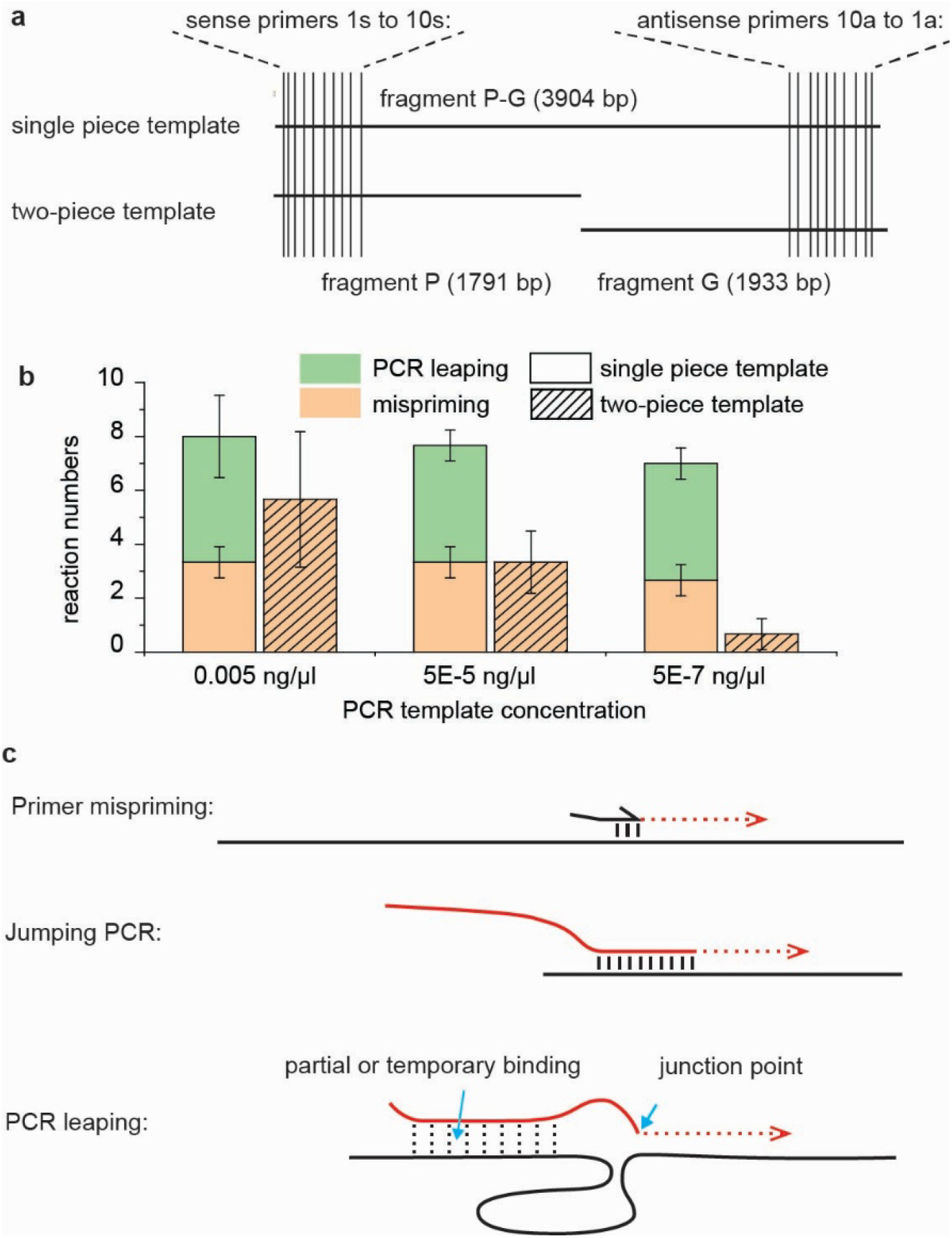
The “same template mechanism” of PCR leaping. **a**, comparison of single piece template and two-piece template. contiguous fragment P-G or a mixture of fragment P and G were used as PCR template. All ten sense primes target sequence P; all antisense primers target sequence G. **b**, summary of the PCR results. Error bars are displaying the ±SD of three replicates. **c**, mechanisms of PCR leaping and other PCR artifacts. Templates are shown as black lines.

Premature extension products are shown as red lines. In mispriming, the primer anneals to an unintended place and gets extended. Because of the high primer concentration, short sequence homology is sufficient. In jumping PCR, the 3’ end of a premature extension product anneals to an unintended place. Long homology with Tm comparable to the PCR reaction temperature is required. In PCR leaping, the 3’ end of a premature extension product can get extended at a wrong place with no homology, because its upstream is partially bonded to or still entwined around the same template during DNA breathing or incomplete renaturation.

To further compare the intra- and inter-molecule models, ten BAC plasmids with different inserts were pooled together and amplified using the universal primers that flank the cloning site. If it is by the “jumping PCR” mechanism, we would expect nine times more intra-molecule products than inter-molecule products. A well-mixed PCR solution was divided into ten aliquots, and all were subjected to the same PCR program. The majority of the products were confirmed to be leaping products from single plasmids with overlaps of 0 to 10 bp, supporting the intramolecular model (supplementary figure 9).

Based on the above observations, we proposed a “same template mechanism” for PCR leaping(Figure 2c). The leaping structure is generated when the 3’ end of a premature extension product approaches a random point of the same template and gets extended. Interaction between the upstream sequence and the template provided stabilization for this to happen, thus PCR leaping can occur without sequence overlap at the leaping points, can outcompete mispriming and occurs only within the same template^30^.

### PCR leaping is overlooked in CRISPR editing validation

CRISPR holds great promise for a wide range of biological and medical applications. However, unintended large deletions at editing sites remain a common concern{Kosicki, 2018 #3854}. Long-range PCR is widely used to detect these unintended deletions and to characterize intended large genomic modifications. Unlike mispriming products, leaping products cannot be filtered out from sequencing data and their junctions mimic the results of Non-Homologous End Joining (NHEJ) and Microhomology-Mediated End Joining (MMEJ), making these artifacts particularly misleading. Here, we reanalyzed the raw data from previous CRISPR editing studies in which long-range PCR and full-length amplicon sequencing were performed. Worryingly, many of these studies did not include negative controls to validate the PCR step. Among the studies that did include negative controls, from which no deletions were expected, PCR leaping ratios are found to be 9.7% for a 5.5 kb amplification{Kosicki, 2018 #3854} and 0.065% for a 4.7 kb amplification{Cullot, 2025 #3855}. In a study that amplified a 9.5 kb area containing two direct homologous repeats of 3.6 kb each{Boutin, 2026 #3856}, only 62.3% are full length products. 2.3% are leaping products with short or zero overlaps. 35.4% are with long overlaps between the repeats, resulted either from PCR leaping or from template switching (supplementary table 1-3). For comparison, unintended deletion frequencies reported across different CRISPR tools, target loci, and cell types range from close to zero to over 20%{Hwang, 2025 #3858}. If the confirmation of editing outcomes rely only on PCR, unintended deletion in CRISPR editing can be substantially overestimated without taking PCR leaping into consideration.

### High-Throughput BAC Library Screening for Biosynthetic Gene Clusters by Leaping PCR

Microbial natural products have long been a major source of antibiotics and other bioactive compounds, but discovery has stalled due to frequent rediscovery of known molecules. Genome analyses reveal a large, untapped reservoir of biosynthetic gene clusters (BGCs); for example, tools like antiSMASH typically identify 30–60 BGCs in an actinobacterial genome, most of which are not linked to any known compound and collectively accounting for around 5 to 15% of the genome (antismash 8.0 reference).

BAC libraries are a classic approach for capturing these BGCs onto expression vectors. One 384-well plate of a library represents around 5.4× coverage of a 10 Mbp genome, enough to contain all the BGCs from a strain. However, existing screening methods are largely limited to colony PCR using BGC-specific primers or degenerate primers targeting conserved sequences^31^. Here we established a high-throughput screening platform based on the PCR leaping phenomenon. Primers flanking the cloning site were added with 4 nt location barcodes at their 5’end. 24 sense primers and 16 antisense primers make up an array of 384 PCR reactions. Colonies from one library microplate were cultured on a LB agar plate and then printed into the 384 PCR reactions with a plastic pin replicator. The PCR results were pooled together and subjected to illumina sequencing (Figure 3a). From the sequencing result, insertion sequences were obtained by mapping the reads against the genome, as leaping products can indicate the two terminals of the insertions. The locations of the corresponding colonies were interpreted from the primer barcode pairs (Figure 3 b and c). Confident results supported by more than 20 sequencing reads were obtained for over 72% of the colonies (supplementary data 1).

**Figure 3.**
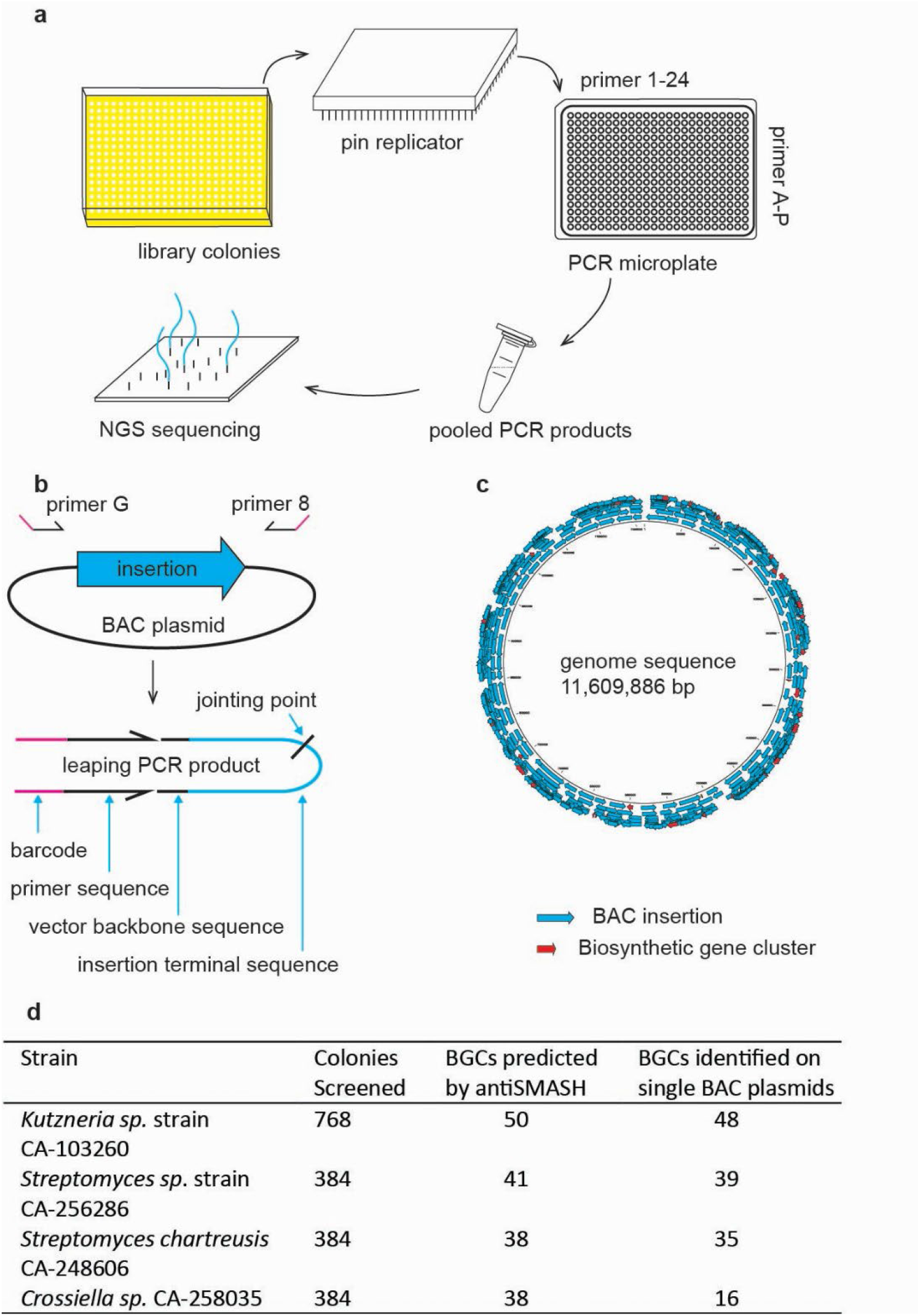
**a**, Leaping PCR based end-paired library analysis. 384 library colonies were transferred from LB agar to multi well PCR plate. The PCR plate is made up from 24 sense primers and 16 antisense primers. The PCR products were pooled together and submitted to NGS sequencing. **b**, BAC plasmid, primer sites and leaping PCR product structure. Primer binding sites flank the cloning site and 6 bp away from the insertions. 4 nt barcodes were added to the 5’end of primers. The terminal sequences of the insertions will be amplified into the PCR leaping products **c**, mapping the leaping PCR product on to the genome. the insertions covering intact BGCs can be identified from the mapping. The corresponding colonies can be found in the library using the barcode pairs. **d**, summery of BAC library screening reuslts.

By this way, we have isolated 138 BGCs from four actinobacterial genomes (Figure 3 d). Full-length sequencing of 20 selected plasmids confirmed that their insertions were all correct (data not shown). The cloned BGCs have contributed to the studies of pteridic acid H and F as new plant hormone ^32^, the discovery of a novel N-acetyl cysteine adduct of antibiotic 3’-O-α-D-forosaminyl-(+)-griseusin A^33^, and the discovery of a new antibacterial cyclic lipopeptide kutzneridine A{Ortiz-López, 2024 #3819}.

## Discussion

PCR has been intensively studied and applied for more than 40 years. Yet, as demonstrated here, our understanding of it can still be improved. Mispriming was a major issue in the early years, when knowledge and tools for primer design were limited. PCR leaping might therefor have been overlooked. Some mystery artifacts were noticed later but not further investigated{Kajander, 1999 #3857}. In regular PCR where the target is not too long or too challenging, leaping products exist in faint smears and do not necessarily interfere with downstream applications, depending on their sensitivity. However, in long range PCR or when stable secondary structures block extension, leaping products can form clear bands or even be the dominant products of the reaction. The leaping product mimics a sequence deletion in both overall and junction structures. This observation has critical impacts on interpreting PCR-based validation strategies in applications like CRISPR editing. On the other hand, PCR leaping can be used intentionally as a simple way of DNA end pairing. Our BAC-mapping platform delivering a high-throughput approach to map BAC-libraries to reference genomes demonstrated high accuracy and was independent of BGC type.

## Materials and Methods

### Materials

OneTaq® Hot Start DNA Polymerases 2X Master Mix and Q5® Hot Start High-Fidelity 2X Master Mix were purchased from NEB. DreamTaq™ Hot Start DNA Polymerases 2X Master Mix, Taq DNA Polymerase and Lambda DNA were purchased from Thermo Scientific. Primers were synthesized by Integrated DNA Technologies with standard desalting. BAC libraries were constructed by Bio S&T Inc. Sanger sequencing service is from Eurofin. Illuminae sequencing service is from was provided by Novogene inc. Munich, Germany, using either the TruSeq PCRfree kit (Illumina inc, San Diego, California, USA), the NEBNext Ultra II DNA Library Prep Kit (Cat No. E7645), or the proprietary Novogene NGS DNA Prep Set kit (Cat No.PT004), all without shearing of the input DNA and without size selection of the final libraries, and the latter two with only 6 PCR cycles. All illumina data is 2×150 nt paired end reads to ensure that both extremities of the leaping PCR product are captured.

DNA fragments, C and P, were amplified from *Corynebacterium variabile* DSM 44702 and *Pseudomonas putida* KT2440 genomes respectively. A combined fragment CP was amplified from a recombinant plasmid with C and P cloned together (supplementary). PCR reactions with 10 different primer pairs were carried out with either fragment CP or a mixture of C and P as templates. All sense primers target the sequence of C, and the all antisense primers target the sequences of P)

### PCR

PCR reaction mixtures were prepared on ice following the manufacturer’s instructions. Regular PCR and 96 well plate PCR were performed on Bio-rad C1000 Touch Thermal Cyclers. 384 well plate PCR were prepared manually or by Echo liquid handler and performed on Veriti™ Dx 384-well Thermal Cycler from Applied Biosystems™.

All PCR programs are listed in supplementary table 4. Primers are listed in supplementary table 5.

### Sequencing of PCR products

Sanger sequencing was performed by Eurofins using the same primers employed for PCR amplification. Illumina sequencing was performed by GENEWIZ.

### Re-examing sequencing data from previous studies

The sequencing data were downloaded from the European Nucleotide Archive and NCBI Sequence Read Archive database. PacBio raw data were first assembled into HiFi reads using the ccs tool. Reads from the negative control sample were extracted based on the NCbarcode sequence. The resulting FASTQ file was mapped to the reference sequence using minimap2, and the alignment was visualized in IGV. Contamination and sequencing artifacts were filtered out manually. Nanopore data were filtered by keeping only the reads that had both primers extended by at least 10 bp using custom Python scripts.

The results were then mapped to the reference sequences and visualized in IGV. Junctions in the PCR leaping products were identified by blast against the reference sequences.

### Data analysis

For analyzing the PCR leaping results, we wrote a shell script which is available at github (https://github.com/tuspjo/LeapingPCR_analysis). The script LeapingPCR_analysis (v.1.0) consists of the following steps. First, the illumina reads are quality and adapter trimmed using AdapterRemoval ^34^(v.2.2.2), with the following settings: --trimns –trimqualities –minlength 100. Second, Adapterremoval is used to demultiplex based on a predefined LeapingPCR primer set, being careful to demultiplex both forward-reverse read pairs, and reverse-forward read pairs, as the direction of sequencing is random. As the number of predefined LeapingPCR indices (384) exceeds the maximum number of indices allowed by Adapter removal, this demultiplexing is run in two batches. Third, the first 25 nt immediately following the spacer TTAGCC is extracted for each read pair. Fourth, the read pairs are mapped with each read individually using bowtie2 (v.2.3.4.1) ^35^ with the following parameters: --end-to-end --no-unal. Fifth, the most common start position for the mapped reads of each barcode is extracted and stored. Sixth, a GFF3 file is built from the indexing name and position in the 384 well plate, the most common start position of each of the pairs of a barcode, and the number of reads which supports the starting positions. Seventh and finally, BACs found to be shorter than 10kb or longer than 500kb are removed, as these are thought to exceed the relevant size range of BACs.

### Confirming the Leaping PCR prediction

To confirm the Leaping PCR results, we performed illumina whole genome sequencing of 20 predicted BACs, using the same kit as mentioned above, and a 2×150 paired end library strategy. The reads were quality and adapter trimmed as above. We mapped the illumina reads to the base genome using Bowtie2 with parameters as above. In all 20 cases, the predicted start and stop positions were correct and the BAC contained the complete sequence between these two positions (SI XXX?)

One*Taq*® Quick-Load® 2X Master Mix with Standard Buffer M0486

Q5® Hot Start High-Fidelity 2X Master Mix M0494 fused with dsdna binding protein^29^

DreamTaq™ Hot Start PCR Master Mix Catalog number: K9011

“OneTaq® DNA Polymerase is an optimized blend of Taq and Deep Vent® DNA polymerases for use with routine and difficult PCR experiments”

## Acknowledgments

This work was funded by grants of the Novo Nordisk Foundation (NNF20CC0035580, NNF25SA0109652, NNF16OC0021746) and by a research grant (36230) from VILLUM FONDEN

