## supplementary figures for "A long-hidden prevalent PCR artifact and its application in DNA library screening"

a.

| reaction | Full amplicon size (bp) | Primer |
| --- | --- | --- |
| 1 | 1970 | PUC1 |
|  |  | PUC2 |
| 2 | 2027 | PUC3 |
|  |  | PUC4 |
| 3 | 1941 | PUC5 |
|  |  | PUC6 |
| 4 | 2278 | PUC7 |
|  |  | PUC8 |
| 5 | 1933 | PUC9 |
|  |  | PUC10 |
| 6 | 2249 | PUC11 |
|  |  | PUC12 |
| 7 | 2102 | PUC13 |
|  |  | PUC14 |
| 8 | 1920 | PUC15 |
|  |  | PUC16 |
| 9 | 2132 | PUC17 |
|  |  | PUC18 |
| 10 | 2149 | PUC19 |
|  |  | PUC20 |
| 11 | 2127 | lambda1 |
|  |  | lambda2 |
| 12 | 2086 | lambda3 |
|  |  | lambda4 |
| 13 | 2205 | lambda5 |
|  |  | lambda6 |
| 14 | 2115 | lambda7 |
|  |  | lambda8 |
| 15 | 2129 | lambda9 |
|  |  | lambda10 |
| 16 | 2292 | lambda11 |
|  |  | lambda12 |
| 17 | 2185 | lambda13 |
|  |  | lambda14 |
| 18 | 2095 | lambda15 |
|  |  | lambda16 |
| 19 | 1948 | lambda17 |
|  |  | lambda18 |
| 20 | 2300 | lambda19 |
|  |  | lambda20 |

b.

c.
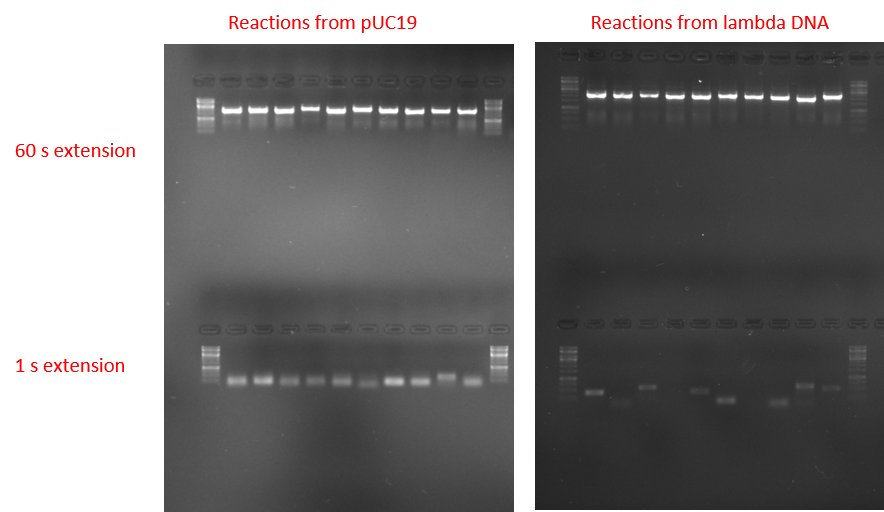

| reaction | Brief of the most abundant reads | Reads and read counts (size) |
| --- | --- | --- |
| 1 | A Leaping product of 126 bp mediated by  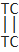  A leaping product of 38+147 bp mediated by 0 bp overlap  A leaping product of 67+219 bp mediated by 0 bp overlap | >A01426:204:HTWM5DSX2:3:1101:11659:1329;size=35  >A01426:204:HTWM5DSX2:3:1101:16297:1282;size=24  >A01426:204:HTWM5DSX2:3:1101:25491:2331;size=21 |
| 2 | A mispriming product of 248 bp from PUC4 mediated by 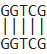  A leaping product of 220 bp mediated by 0 bp overlap  A leaping product of 223 bp mediated by 0 bp overlap | >A01426:204:HTWM5DSX2:3:1101:7455:1689;size=72  >A01426:204:HTWM5DSX2:3:1101:9146:2112;size=28  >A01426:204:HTWM5DSX2:3:1101:18087:1689;size=25 |
| 3 | A leaping product of 180 bp mediated by 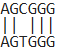  A leaping product of 146 bp mediated by 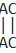  A leaping product of 211 bp mediated by 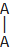 | >A01426:204:HTWM5DSX2:3:1101:20419:4507;size=39  >A01426:204:HTWM5DSX2:3:1101:22263:1939;size=26  >A01426:204:HTWM5DSX2:3:1101:11514:10786;size=20 |
| 4 | A leaping product of 162 bp mediated by 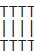  A leaping product of 158 bp mediated by 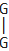  A leaping product of 299 bp mediated by 0 bp | >A01426:204:HTWM5DSX2:3:1101:12210:1470;size=58  >A01426:204:HTWM5DSX2:3:1101:28447:4351;size=16  >A01426:204:HTWM5DSX2:3:1101:23999:12618;size=14 |
| 5 | A leaping product of 139 bp mediated by 0 bp | >A01426:204:HTWM5DSX2:3:1101:4119:1235;size=116 |
| 6 | A leaping product of 444 bp mediated by 0 bp | >A01426:204:HTWM5DSX2:3:1101:4508:3537;size=3 |
| 7 | A leaping product of 83 bp mediated by 0 bp  A leaping product of 165 bp mediated by 0 bp | >A01426:204:HTWM5DSX2:3:1101:3513:1063;size=217  >A01426:204:HTWM5DSX2:3:1101:14027:2268;size=21 |
| 8 | A leaping product of 144 bp mediated by 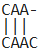  A leaping product of 136 bp mediated by 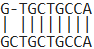 | >A01426:204:HTWM5DSX2:3:1101:20112:1877;size=119  >A01426:204:HTWM5DSX2:3:1101:18276:1955;size=16 |
| 9 | A leaping product of 297 bp mediated by 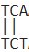  A leaping product of 366 bp mediated by  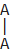 | >A01426:204:HTWM5DSX2:3:1101:12472:16705;size=4  >A01426:204:HTWM5DSX2:3:1102:14579:22576;size=3 |
| 10 | A leaping product of 137 bp mediated by 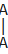  A leaping product of bp mediated by 187 bp mediated by 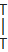 | >A01426:204:HTWM5DSX2:3:1101:21450:1031;size=98  >A01426:204:HTWM5DSX2:3:1101:9977:3176;size=16 |
| 11 | A Mispriming product of 411 bp from lambda2 mediated by 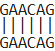 | >A01426:204:HTWM5DSX2:3:1101:29894:1501;size=73 |
| 12 | A Mispriming product of 103 bp from both lambda3 and lambda4 mediated by  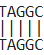 and 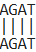 | >A01426:204:HTWM5DSX2:3:1101:15049:1501;size=43 |
| 13 | A leaping product of 577 bp mediated by 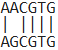 | >A01426:204:HTWM5DSX2:3:1101:27380:15655;size=22 |
| 14 | A leaping product of 430 bp mediated by 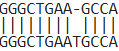 | >A01426:204:HTWM5DSX2:3:1102:6397:16141;size=1 |
| 15 | A Mispriming product of 426 bp from lambda9 mediated by 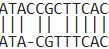 | >A01426:204:HTWM5DSX2:3:1101:26594:11819;size=29 |
| 16 | A leaping product of 163 bp mediated by 0 bp overlap | >A01426:204:HTWM5DSX2:3:1101:23267:1986;size=59 |
| 17 | No corresponding read (also no band on the gel) |  |
| 18 | A Mispriming product of 79 bp from lambda15 mediated by  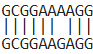 | >A01426:204:HTWM5DSX2:3:1101:10031:22404;size=1 |
| 19 | A leaping product of 607 bp mediated by 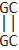 | >A01426:204:HTWM5DSX2:3:1101:24885:1282;size=10 |
| 20 | A Mispriming product of 517 bp from both lambda19 and lambda20 mediated by  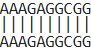 and 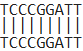 | >A01426:204:HTWM5DSX2:3:1101:4200:4194;size=7 |

Figure S1. **Enrichment of artifact products by reducing extension time.** The reaction mixtures were the same as in Figure 1. By reducing the extension time from 60 s to 1 s, full-length products disappeared. The artifact products were pooled and subjected to Illumina sequencing. **a,** reactions and target sizes. **b,** the gel image. **c,** summary of the sequencing results.

a.

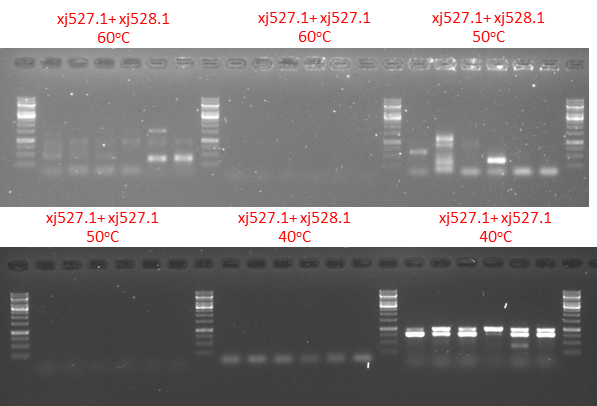

b.

| **Access number** | **reaction** | **repeat number** | **sequencing primer** |
| --- | --- | --- | --- |
| EF72285197 | Xj527.1+ XJ528.1 60oC | 1 | xj527.1 |
| EF72285198 | Xj527.1+ XJ528.1 60oC | 2 | xj527.1 |
| EF72285199 | Xj527.1+ XJ528.1 60oC | 3 | xj527.1 |
| EF72285200 | Xj527.1+ XJ528.1 60oC | 4 | xj527.1 |
| EF72285201 | Xj527.1+ XJ528.1 60oC | 5 | xj527.1 |
| EF72285202 | Xj527.1+ XJ528.1 60oC | 6 | xj527.1 |
| EF72285203 | Xj527.1+ XJ528.1 60oC | 1 | xj528.1 |
| EF72285204 | Xj527.1+ XJ528.1 60oC | 2 | xj528.1 |
| EF72285205 | Xj527.1+ XJ528.1 60oC | 3 | xj528.1 |
| EF72285206 | Xj527.1+ XJ528.1 60oC | 4 | xj528.1 |
| EF72285207 | Xj527.1+ XJ528.1 60oC | 5 | xj528.1 |
| EF72285208 | Xj527.1+ XJ528.1 60oC | 6 | xj528.1 |
| EF72285209 | Xj527.1+ XJ528.1 50oC | 1 | xj527.1 |
| EF72285210 | Xj527.1+ XJ528.1 50oC | 2 | xj527.1 |
| EF72285211 | Xj527.1+ XJ528.1 50oC | 3 | xj527.1 |
| EF72285212 | Xj527.1+ XJ528.1 50oC | 4 | xj527.1 |
| EF72285213 | Xj527.1+ XJ528.1 50oC | 5 | xj527.1 |
| EF72285214 | Xj527.1+ XJ528.1 50oC | 6 | xj527.1 |
| EF72285215 | Xj527.1+ XJ528.1 50oC | 1 | xj528.1 |
| EF72285216 | Xj527.1+ XJ528.1 50oC | 2 | xj528.1 |
| EF72285217 | Xj527.1+ XJ528.1 50oC | 3 | xj528.1 |
| EF72285218 | Xj527.1+ XJ528.1 50oC | 4 | xj528.1 |
| EF72285219 | Xj527.1+ XJ528.1 50oC | 5 | xj528.1 |
| EF72285220 | Xj527.1+ XJ528.1 50oC | 6 | xj528.1 |
| EF72285221 | Xj527.1+ Xj527.1 40oC | 1 | xl29 |
| EF72285222 | Xj527.1+ Xj527.1 40oC | 2 | xl29 |
| EF72285223 | Xj527.1+ Xj527.1 40oC | 3 | xl29 |
| EF72285224 | Xj527.1+ Xj527.1 40oC | 4 | xl29 |
| EF72285225 | Xj527.1+ Xj527.1 40oC | 5 | xl29 |
| EF72285226 | Xj527.1+ Xj527.1 40oC | 6 | xl29 |
| EF72285227 | Xj527.1+ Xj527.1 40oC | 1 | xj30 |
| EF72285228 | Xj527.1+ Xj527.1 40oC | 2 | xj30 |
| EF72285229 | Xj527.1+ Xj527.1 40oC | 3 | xj30 |
| EF72285230 | Xj527.1+ Xj527.1 40oC | 4 | xj30 |
| EF72285231 | Xj527.1+ Xj527.1 40oC | 5 | xj30 |
| EF72285232 | Xj527.1+ Xj527.1 40oC | 6 | xj30 |

c.

| **sample** | **repeat number** | **Summary of the sequencing result** |
| --- | --- | --- |
| Xj527.1+ XJ528.1 60oC | 1 | Not able to tell |
| Xj527.1+ XJ528.1 60oC | 2 | Not able to tell |
| Xj527.1+ XJ528.1 60oC | 3 | Not able to tell |
| Xj527.1+ XJ528.1 60oC | 4 | Not able to tell |
| Xj527.1+ XJ528.1 60oC | 5 | Leaping pcr sequencing did reach the leaping point |
| Xj527.1+ XJ528.1 60oC | 6 | Leaping pcr mediated by 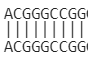 |
| Xj527.1+ XJ528.1 50oC | 1 | Leaping pcr, mediated by 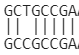 |
| Xj527.1+ XJ528.1 50oC | 2 | Not able to tell |
| Xj527.1+ XJ528.1 50oC | 3 | Not able to tell |
| Xj527.1+ XJ528.1 50oC | 4 | Leaping pcr mediated by 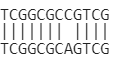 |
| Xj527.1+ XJ528.1 50oC | 5 | Not able to tell |
| Xj527.1+ XJ528.1 50oC | 6 | Leaping pcr mediated by 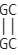 |
| Xj527.1+ Xj527.1 40oC | 1 | Mispriming on E.coli dna by 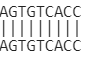 |
| Xj527.1+ Xj527.1 40oC | 2 | Not able to tell |
| Xj527.1+ Xj527.1 40oC | 3 | Mispriming on E.coli dna by 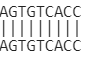 |
| Xj527.1+ Xj527.1 40oC | 4 | Not able to tell |
| Xj527.1+ Xj527.1 40oC | 5 | Not able to tell |
| Xj527.1+ Xj527.1 40oC | 6 | Not able to tell |

**Figure S2.** **Comparison between single primer PCR and PCR leaping with the same pR12-13BACP4e1 template and different touchdown final annealing temperatures.** Primer Xj527.1	and XJ528.1 flank the cloning site. Insertion is a 223 kb fragment (sequence 1731079-1953942 of *Kutzneria* sp. CA-103260 genome NZ_CP073318.1, average GC content of 72%). Each reaction was repeated 6 times. OneTaq® Quick-Load® 2X Master was used for all reactions. The reaction mixtures are prepared according to the product instruction. Primers are used at a final concentration of 0.2 µM. Template is used at a final concentration of 0.1 ng/μl. The PCR programs with different final touchdown annealing temperatures OnetaqTD60, OnetaqTD50 and can be found in supplementary table 1. Primer sequences can be found in supplementary table 2. a, gel photo of the pcr products. b, access numbers of the sanger sequencing result. c, summary of the sequencing results. Sanger sequencing data can be found in supplementary data 1 11108861865-1 and 11108869947-1.

a.

| **Sample number** | **primers** | **pR12-13BACP4e1 Template (0,5ul used in 20 ul reaction)** |
| --- | --- | --- |
| 1 | Xj527.1+XJ528.1 | 0.1 ng/ul Plasmid dna |
| 2 | Xj527.1+XJ462 | 0.1 ng/ul Plasmid dna |
| 3 | XJ528.1+XJ462 | 0.1 ng/ul Plasmid dna |
| 4 | Xj527.1+XJ528.1 | E.coli colony |
| 5 | Xj527.1+XJ462 | E.coli colony |
| 6 | XJ528.1+XJ462 | E.coli colony |
| 7 | Xj527.1+XJ528.1 | E.coli colony 99oC in 100 ul water for 10min. |
| 8 | Xj527.1+XJ462 | E.coli colony 99oC in 100 ul water for 10min. |
| 9 | XJ528.1+XJ462 | E.coli colony 99oC in 100 ul water for 10min. |

b.

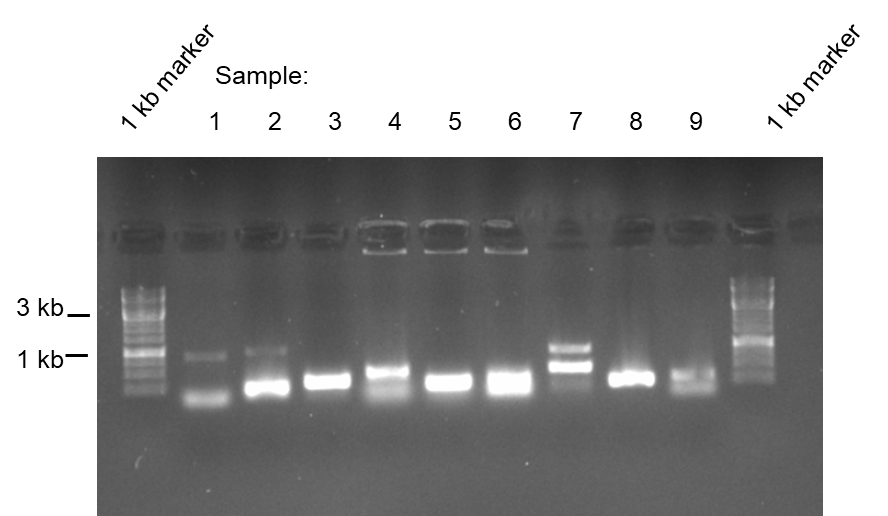

c.

| **Sample number** | Xj527.1 | XJ528.1 | XJ462 |
| --- | --- | --- | --- |
| 1 | EF30321583 | EF30321594 |  |
| 2 | EF30321585 |  | EF30321595 |
| 3 |  | EF30321596 | EF30321586 |
| 4 | EF30321587 | EF30321597 |  |
| 5 | EF30321588 |  | EF30321548 |
| 6 |  | EF30321599 | EF30321589 |
| 7 | EF30321590 | EF30321600 |  |
| 8 | EF30321592 |  | EF30321602 |
| 9 |  | EF30321603 | EF30321593 |

d.

| **Sample number** | **Summary of sequencing result** |
| --- | --- |
| 1 | PCR leaping |
| 2 | Primer mispriming by XJ462 |
| 3 | Primer mispriming by XJ462 |
| 4 | PCR leaping |
| 5 | Primer mispriming by XJ462 |
| 6 | Primer mispriming by XJ462 |
| 7 | PCR leaping |
| 8 | Primer mispriming by XJ462 |
| 9 | Primer mispriming by XJ462 |

**Figure S3. PCR products from different template preparations.** BAC insertion is a 223 kb fragment (sequence 1731079-1953942 of *Kutzneria* sp. CA-103260 genome NZ_CP073318.1, average GC content of 72%). Primer Xj527.1	and XJ528.1 flank the cloning site. Xj 462 is a random primer that has not binding site on the plasmid pR12-13BACP4e1. OneTaq® Quick-Load® 2X Master Mix with Standard Buffer and program OnetaqTD40 (supplementary table 1) were used for all reactions. The reaction mixtures are prepared according to the product instruction. Primers are used at a final concentration of 0.2 µM. a, primers and template used for each sample. b, gel photo of the pcr products. c, access number of the sanger sequencing result for each product. d, summary of the sequencing results. Primer sequences can be found in supplementary table 2. Sanger sequencing data can be found in supplementary data 1 11106506667-1.

a.

| **Sample number** | **primers** | **Template (0,5ul LB overnight culture used in 20 ul reaction)** |
| --- | --- | --- |
| 1 | Xj527.2+XJ528.2 | #QUO5687 plate 1 5B |
| 2 | Xj527.2+XJ528.2 | #QUO5687 plate3 3L |
| 3 | Xj527.2+XJ528.2 | #QUO5687 plate3 4D |
| 4 | Xj527.2+XJ528.2 | #QUO5687 plate 7 11J |
| 5 | Xj527.2+XJ528.2 | #QUO5697 plate1 3A |
| 6 | Xj527.2+XJ528.2 | #QUO5697 plate3 8C |
| 7 | Xj527.2+XJ528.2 | #QUO5697 plate 4 1E |
| Empty control | Xj527.2+XJ528.2 | Water |

b.

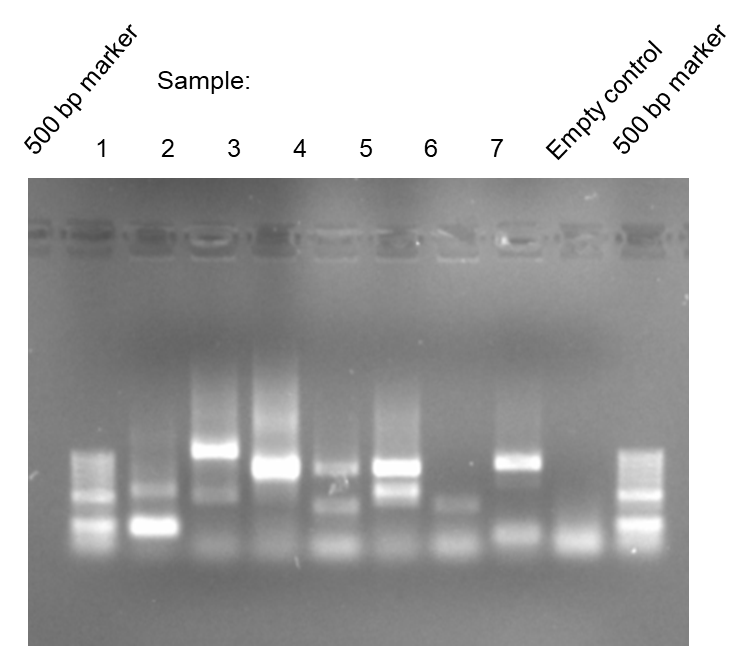

c.

| Samples from Q5 | Sense xj527.1 | Antisense xj528.1 |
| --- | --- | --- |
| #QUO5687 plate 1 5B | EF30321550 | EF30321582 |
| #QUO5687 plate3 3L | EF30321573 | EF01115418 |
| #QUO5687 plate3 4D | EF30321576 | EF01115419 |
| #QUO5687 plate 7 11J | EF30321577 | EF01115420 |
| #QUO5697 plate1 3A | EF30321578 | EF30321694 |
| #QUO5697 plate3 8C | EF30321579 | EF30320595 |
| #QUO5697 plate 4 1E | EF30321580 | EF30321695 |

d.

| **Sample number** | **Summary of sequencing result** |
| --- | --- |
| 1 | Mispriming by xj528.2 |
| 2 | PCR leaping mediated by  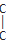 |
| 3 | PCR leaping |
| 4 | PCR Leaping mediated by  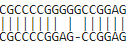 |
| 5 | Not able to tell |
| 6 | Mispriming by xj528.2 |
| 7 | PCR leaping mediated by  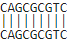 |

**Figure S4. PCR products from proofreading DNA polymerase.** Q5 Hot Start High-Fidelity 2X Master Mix was used. The reaction mixtures was prepared according to the product instruction. Primers were used at a final concentration of 0.5 µM. 0.5 μl *E.coli* LB cultures of seven colonies from the BAC library #QUO5687 (made from gDNA of Streptomyces sp. NBC_00104 CP108215.1) were used as templates in 20 μl reactions. The PCR programs Q5TD40 used here can be found in supplementary table 1. Primer sequences can be found in supplementary table 2. a, primers and template used for each sample. b, gel photo of the PCR products. c, access number of the sanger sequencing result for each product. d, summary of the sequencing results. Sanger sequencing data can be found in supplementary data 1 11106562108-1.

a.

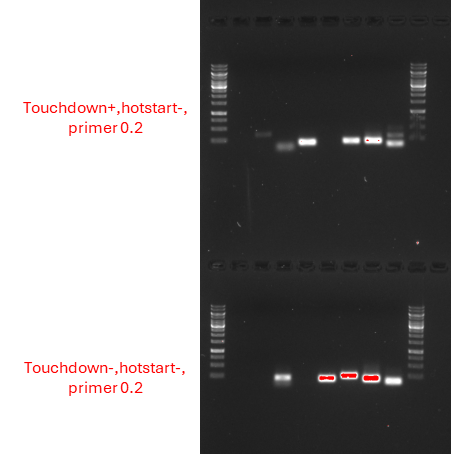

b.

|  | **repeat number** | **From primer** PUC13 | **From primer** PUC14 |
| --- | --- | --- | --- |
| touchdown+ hotstart- | 1 | EF70920142 | EF70919785 |
|  | 2 | EF70920140 | EF70911263 |
|  | 3 | EF70919811 | EF70919787 |
|  | 4 | EF70919812 | EF70919788 |
|  | 5 | EF70919821 | EF70919797 |
|  | 6 | EF70919822 | EF70920208 |
|  | 7 | EF70919823 | EF70919799 |
|  | 8 | EF70919824 | EF70919800 |
| Touchdown- hotstart- | 1 | EF70919833 | EF70911286 |
|  | 2 | EF70919834 | EF70911310 |
|  | 3 | EF70911251 | EF70911299 |
|  | 4 | EF70919836 | EF70911262 |
|  | 5 | EF70911298 | EF70919829 |
|  | 6 | EF70911311 | EF70919830 |
|  | 7 | EF70911287 | EF70919831 |
|  | 8 | EF70919840 | EF70919832 |

c.

|  | **repeat** | **Summary of the sequencing result** |
| --- | --- | --- |
| touchdown+ hotstart- | 1 | Sequencing failed |
|  | 2 | Leaping pcr mediated by  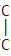 |
|  | 3 | Primer dimer |
|  | 4 | Leaping pcr  0 overlap |
|  | 5 | Sequencing failed |
|  | 6 | Leaping pcr mediated by  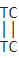 |
|  | 7 | Leaping pcr mediated by  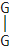 |
|  | 8 | Leaping pcr mediated by  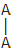  And  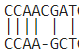 |
| Touchdown- hotstart- | 1 | Sequencing failed |
|  | 2 | Sequencing failed |
|  | 3 | Leaping pcr mediated by  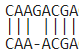 |
|  | 4 | Sequencing failed |
|  | 5 | Leaping pcr  No overlap |
|  | 6 | Leaping pcr  No overlap |
|  | 7 | Leaping pcr mediated by  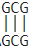 |
|  | 8 | First a premature product of GACTGCTGCAGTGGCGATAAGTCGTGTCTTAC  CGGGTTGGACTCAAGACGATAGTTACCGGATAAGGCG  Misbinded with the middle part of antisense primer by overlap  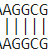  Then the extended product misbinded with the template (  Leaping pcr)  By overap  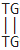 |

**Figure S5. PCR products from different programs without hot-start and touchdown.** Reactions were prepared with Taq DNA Polymerase (thermofisher EP0402, it is without hotstart function) according to the product instruction. pUC19 DNA was used as template at a final concentration of 1 pg/µl. Primer PUC13 and PUC14 were used at a final concentration of 0.2 µM. The same reaction was repeated 8 times. The program TaqTD50 (touchdown program) and Taq50 (non touchdown program) used here can be found supplementary table 1. Primer sequences can be found in supplementary table 2. a, gel photo of the PCR products. b, access number of the sanger sequencing result for each product. c, summary of the sequencing results. Sanger sequencing data can be found in supplementary data 1 11108959574-1.

a.

| **Sample number** | **primers** | **Template (0,5ul LB overnight culture used in 20 ul reaction)** |
| --- | --- | --- |
| 1 | Xj527.7+XJ528.7 | #QUO5687 plate 1 5B |
| 2 | Xj527.2+XJ528.2 | #QUO5687 plate3 3L |
| 3 | Xj527.2+XJ528.2 | #QUO5687 plate3 4D |
| 4 | Xj527.2+XJ528.2 | #QUO5687 plate 7 11J |
| 5 | Xj527.2+XJ528.2 | #QUO5697 plate1 3A |
| 6 | Xj527.2+XJ528.2 | #QUO5697 plate3 8C |
| 7 | Xj527.2+XJ528.2 | #QUO5697 plate 4 1E |
| Empty control | Xj527.2+XJ528.2 | water |

b.

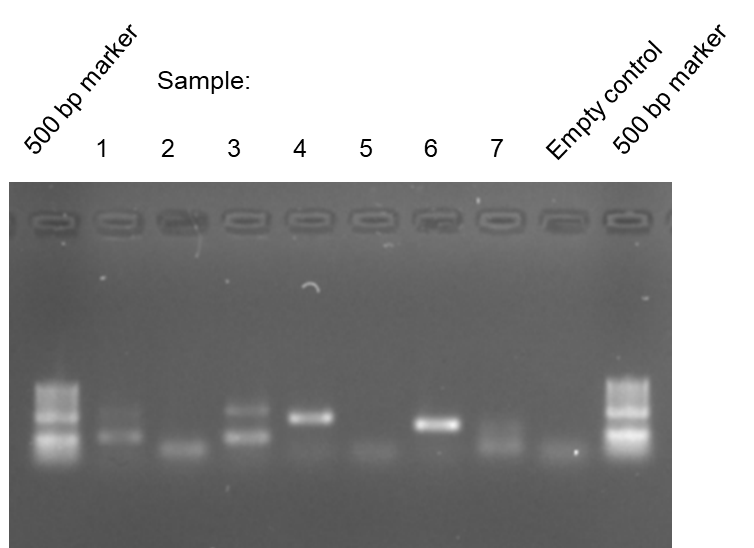

c.

|  | Samples | Sense xj527.1 | Antisense xj528.1 |
| --- | --- | --- | --- |
| Amplified from xj527.7+xj528.7 | #QUO5687 plate 1 5B | EF30321557 | EF30321564 |
|  | #QUO5687 plate3 3L | EF30321558 | EF30321565 |
|  | #QUO5687 plate3 4D | EF30321559 | EF30321566 |
|  | #QUO5687 plate 7 11J | EF30321560 | EF30321567 |
|  | #QUO5697 plate1 3A | EF30321561 | EF30321568 |
|  | #QUO5697 plate3 8C | EF30321562 | EF30321569 |
|  | #QUO5697 plate 4 1E | EF30321563 | EF30321570 |

d.

| **Sample number** | **Summary of sequencing result** |
| --- | --- |
| 1 | PCR Leaping, sequencing did not reach the leaping point |
| 2 | Poor sequencing result |
| 3 | PCR leaping mediated by  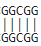 |
| 4 | Mispriming by xj528.7 |
| 5 | Poor sequencing result |
| 6 | PCR leaping mediated by  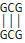 |
| 7 | PCR Leaping, sequencing did not reach the leaping point |

**Figure S6. PCR products from primers with hairpin structure protection.** Reactions were prepared with OneTaq® Quick-Load® 2X Master Mix with Standard Buffer according to the product instruction and supplemented with 1 M betaine(final concentration). 0.5 ul liquid cultures were used as template. Primers xj527.7 and xj528.7 are used at a final concentration of 0.2 µM. The program OnetaqTD40b used here can be found supplementary table 1. Primer sequences can be found in supplementary table 2. a, primers and template used for each sample. b, gel photo of the PCR products. c, access number of the sanger sequencing result for each product. d, summary of the sequencing results. Sanger sequencing data can be found in supplementary data 1 11106562108-1.

a.

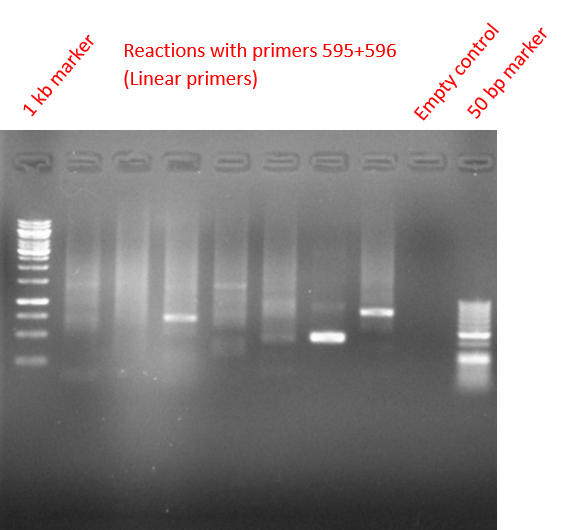

b.

| colony | access number of the sanger sequencing result | brief of the sequencing result |
| --- | --- | --- |
| Colony 1 | >EF31178875_EF31178875  359820 to 360077 of Vibrio natriegens chromosome 1, CP009977.1  >EF31178900_EF31178900  E.coli K12 sequence | Contaminated with E.coli sequence |
| Colony 2 | >EF31178876_EF31178876  1087942 to 1088693 of Vibrio natriegens chromosome 2, CP009978.1  >EF31178902_EF31178902  1072530 to 1073216 of Vibrio natriegens chromosome 2, CP009978.1 | Leaping pcr. The insertion size is 16163 bp |
| Colony 3 | >EF31178877_EF31178877  1078475 to 1078994of Vibrio natriegens chromosome 1, CP009977.1  >EF31178903_EF31178903  1093457 to 1094087 of Vibrio natriegens chromosome 1, CP009977.1 | Leaping pcr. The insertion size is 15.612 |
| Colony 4 | >EF31178878_EF31178878  1425744 to 1426060 of Vibrio natriegens chromosome 2, CP009978.1  >EF31178904_EF31178904  1399516 to 1399748 of Vibrio natriegens chromosome 2, CP009978.1 | Leaping pcr. The insertion size is 26.544 bp |
| Colony 5 | >EF31178879_EF31178879  2706048 to 2706275 of Vibrio natriegens chromosome 1, CP009977.1  >EF31178905_EF31178905  2697551 to 2697891 of Vibrio natriegens chromosome 1, CP009977.1 | Leaping pcr. The insertion size is 8.724 bp |
| Colony 6 | >EF31178880_EF31178880  460493 to 460588 of Vibrio natriegens chromosome 1, CP009977.1  >EF31178906_EF31178906  469674 to 469945 of Vibrio natriegens chromosome 1, CP009977.1 | Leaping pcr. The insertion size is 9.452 bp  Mediated by AAGCTT |
| Colony 7 | >EF31178882_EF31178882  165009 to 165201 of Vibrio natriegens chromosome 2, CP009978.1  >EF31178907_EF31178907  151769 to 152482 of Vibrio natriegens chromosome 2, CP009978.1 | Leaping pcr. The insertion size is 13.432 bp |
| Empty control | >EF31178908_EF31178908  >EF31178884_EF31178884 |  |

**Figure S7. Identification of DNA library insertions (45% GC content) by PCR leaping**. Seven colonies from the DNA library of *Vibrio natriegens* DSM 759 genomic DNA were amplified using primers xj595 and xj596, which flank the cloning site. Reactions were prepared with DreamTaq 2X Master Mix with Standard Buffer according to the manufacturer's instructions. Primers were used at a final concentration of 0.1 µM. The thermal cycling program (DreamTaqTD20) is described in Supplementary Table 1, and primer sequences are provided in Supplementary Table 2. **(a)** Agarose gel image of the PCR products. **(b)** Summary of Sanger sequencing results. Raw Sanger sequencing data are available in Supplementary Data. Sanger sequencing data can be found in supplementary data 11106792761-1

a.

| reaction | Target size | Primer |
| --- | --- | --- |
| 1 | 3837 | 1s |
|  |  | 1a |
| 2 | 3781 | 2s |
|  |  | 2a |
| 3 | 3694 | 3s |
|  |  | 3a |
| 4 | 3576 | 4s |
|  |  | 4a |
| 5 | 3482 | 5s |
|  |  | 5a |
| 6 | 3375 | 6s |
|  |  | 6a |
| 7 | 3275 | 7s |
|  |  | 7a |
| 8 | 3189 | 8s |
|  |  | 8a |
| 9 | 3074 | 9s |
|  |  | 9a |
| 10 | 2972 | 10s |
|  |  | 10a |

b.

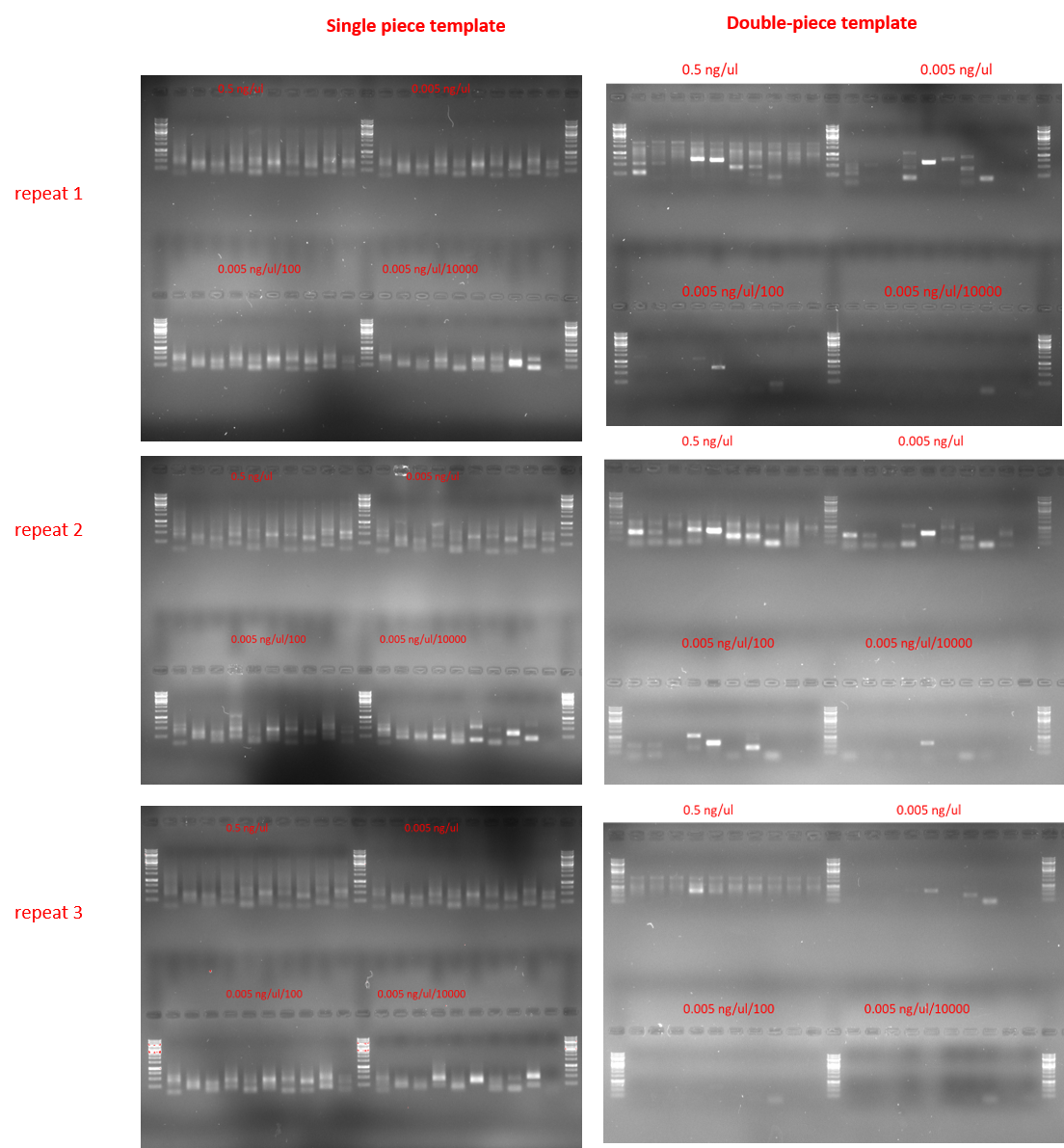

**Figure S8. Comparison of single piece template and double-piece template.** a. reactions and target sizes. b, Gel images of the PCR products in figure 2a and b. Reactions were prepared with DreamTaq 2X Master Mix with Standard Buffer according to the manufacturer's instructions. Primers were used at a final concentration of 0.1 µM. The thermal cycling program (DreamTaqTD50) is described in Supplementary Table 1, and primer sequences are provided in Supplementary Table 2.

a.

| BAC plasmids | Brief* |
| --- | --- |
| pR12-13BACP4e1 | Covering antismash region 12 and 13 |
| pR40BACP2n11 | Covering antismash region 40 |
| pR25BACP1p3 | Covering antismash region 25 |
| pR46BACP1c11 | Covering antismash region 46 |
| pR27BACP1n6 | Covering antismash region 27 |
| pR22BACP2c10 | Covering antismash region 22 |
| pR42BACP1l7 | Covering antismash region 42 |
| pR16BACP2a4 | Covering antismash region 16 |
| pR4BACP2a12 | Covering antismash region 4 |
| pR6BACP1e13 | Covering antismash region 6 |

b.

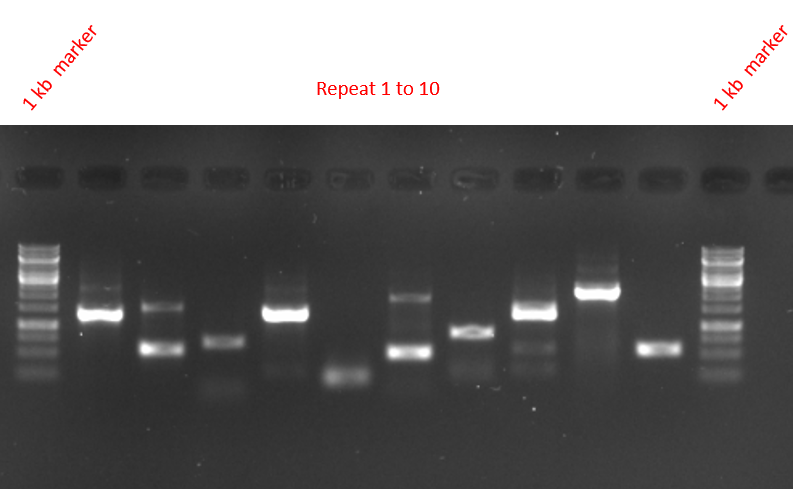

c.

| **Sample name** | **Sequenced by** Xj527.1 | **Sequenced by** XJ528.1 |
| --- | --- | --- |
| mixture aliquot 1 | EF31178959 | EF31315889 |
| mixture aliquot 2 | EF31178961 | EF31315890 |
| mixture aliquot 3 | EF31178963 | EF31315891 |
| mixture aliquot 4 | EF31178964 | EF31315892 |
| mixture aliquot 5 | EF31178965 | EF31315894 |
| mixture aliquot 6 | EF31178966 | EF31315895 |
| mixture aliquot 7 | EF31178968 | EF31315896 |
| mixture aliquot 8 | EF31178969 | EF31315897 |
| mixture aliquot 9 | EF31178970 | EF31315899 |
| mixture aliquot 10 | EF31178971 | EF31315900 |

d.

| Sample name | PCR product | junction |
| --- | --- | --- |
| mixture aliquot 1 | Leaping pcr product from pR40BACP2n11 | 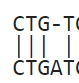 |
| mixture aliquot 2 | Leaping pcr product from pR27BACP1n6 | 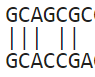 |
| mixture aliquot 3 | Leaping pcr product from pR10BACXX | No overlap |
| mixture aliquot 4 | Leaping pcr product from pR22BACP2c10 | No overlap |
| mixture aliquot 5 | Leaping pcr product from pR25BACP1p3 | 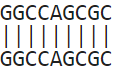  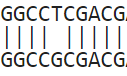 |
| mixture aliquot 6 | Leaping pcr product from pR25BACP1p3 and a leaping pcr product from pR22BACP2c10 | 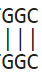  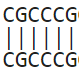 |
| mixture aliquot 7 | Leaping pcr product from pR27BACP1n6 | 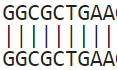 |
| mixture aliquot 8 | Leaping pcr product from pR25BACP1p3 | 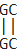 |
| mixture aliquot 9 | Overlap extension product between  pR40BACP2n11+ pR22BACP2c10 | 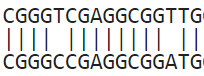 |
| mixture aliquot 10 | Leaping pcr product from pR27BACP1n6 | 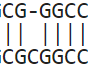 |

**Figure S9 PCR products from mixed BAC plasmids.** a, BAC plasmids used. All plasmid preparations were adjusted to 10ng/ul and then pooled together and used as template. The well-mixed PCR solution was divided into ten aliquots, and all were subjected to the same PCR program DreamtaqTD66. Primer Xj527.1 and XJ528.1 flank the cloning site. Primer sequences can be found in supplementary table 2. b, gel photo of the PCR products. c, access number of the sanger sequencing result for each product. d, summary of the sequencing results. Sanger sequencing data can be found in supplementary data 1 11107008303-1.
