## supplementary tables for "A long-hidden prevalent PCR artifact and its application in DNA library screening"

**Table S1**

**PCR leaping products from previous CRISPR editing study 1. Repair of double-strand breaks induced by CRISPR–Cas9 leads to large deletions and complex rearrangements.**

The PacBio raw data (ERR2512074, ERR2512075, and ERR2512076) were downloaded from the European Nucleotide Archive. The BAM files were first assembled into HiFi reads using the command: ccs subreads.x.bam subreads.x.ccs.bam --minPasses 3 --minPredictedAccuracy 0.9. Reads from the negative control in Figure 2a were extracted using the barcode TCAGACGATGCGTCAT with the command: lima --ccs --same subreads.x.ccs.bam NCbarcode.fasta NCx.bam. The three resulting files were combined using samtools merge negative_control_all.bam NC1.bam NC2.bam NC3.bam. The combined file was converted to FASTQ with samtools fastq negative_control_all.bam > negative_control_all.fastq. The reads were mapped to the reference sequence using minimap2 -t 8 -ax map-hifi pigAamplicon.fasta negative_control_all.fastq | samtools sort -o aligned_negative.bam - and indexed with samtools index aligned_negative.bam. The alignment was then viewed in IGV. Within 350 mapped reads, we found the following 34 leaping products.

| Read | junction | Deletion size (bp) |
| --- | --- | --- |
| m161203_032609_00127_c101108862550000001823246505011795_s1_p0/117658/ccs | 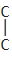 | 2728 |
| m161203_032609_00127_c101108862550000001823246505011795_s1_p0/78603/ccs | 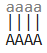 | 1785 |
| m161203_032609_00127_c101108862550000001823246505011795_s1_p0/32244/ccs | 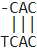 | 3498 |
| m161203_032609_00127_c101108862550000001823246505011795_s1_p0/2444/ccs | 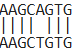 | 3627 |
| m161203_032609_00127_c101108862550000001823246505011795_s1_p0/108002/ccs | 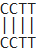 | 2082 |
| m161203_032609_00127_c101108862550000001823246505011795_s1_p0/125602/ccs | 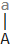 | 1296 |
| m161203_032609_00127_c101108862550000001823246505011795_s1_p0/148671/cc | 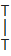 | 1553 |
| m161203_032609_00127_c101108862550000001823246505011795_s1_p0/71042/ccs | 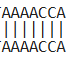 | 1580 |
| m161203_032609_00127_c101108862550000001823246505011795_s1_p0/137975/ccs | 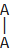 | 3684 |
| m161203_032609_00127_c101108862550000001823246505011795_s1_p0/67653/ccs | 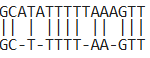 | 2794 |
| m161203_032609_00127_c101108862550000001823246505011795_s1_p0/153240/ccs | No overlap | 1004 |
| m161203_032609_00127_c101108862550000001823246505011795_s1_p0/153240/ccs | 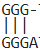 | 849 |
| m161203_032609_00127_c101108862550000001823246505011795_s1_p0/42706/ccs | 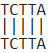 | 3792 |
| m161203_032609_00127_c101108862550000001823246505011795_s1_p0/76807/ccs | 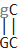 | 3130 |
| m161203_032609_00127_c101108862550000001823246505011795_s1_p0/81613/ccs | 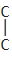 | 2728 |
| m161203_032609_00127_c101108862550000001823246505011795_s1_p0/157225/ccs | 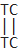 | 1925 |
| m161203_032609_00127_c101108862550000001823246505011795_s1_p0/128876/ccs | 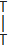 | 3126 |
| m161203_032609_00127_c101108862550000001823246505011795_s1_p0/128876/ccs | 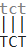 | 309 |
| @m161203_032609_00127_c101108862550000001823246505011795_s1_p0/1293/ccs  m161203_032609_00127_c101108862550000001823246505011795_s1_p0/1293/ccs | 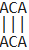 | 852 |
| m161203_032609_00127_c101108862550000001823246505011795_s1_p0/78290/ccs | No overlap | 300 |
| m161203_032609_00127_c101108862550000001823246505011795_s1_p0/65265/cc | 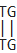 | 1602 |
| m161203_032609_00127_c101108862550000001823246505011795_s1_p0/162177/ccs | 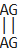 | 2564 |
| m161203_032609_00127_c101108862550000001823246505011795_s1_p0/85164/ccs | No overlap | 4446 |
| m161203_032609_00127_c101108862550000001823246505011795_s1_p0/78865/ccs | No overlap | 4446 |
| m161203_032609_00127_c101108862550000001823246505011795_s1_p0/46212/ccs | No overlap | 4446 |
| m161203_032609_00127_c101108862550000001823246505011795_s1_p0/161603/ccs | 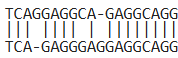 | 4521 |
| m161203_032609_00127_c101108862550000001823246505011795_s1_p0/130197/ccs | 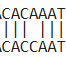 | 4738 |
| m161203_032609_00127_c101108862550000001823246505011795_s1_p0/8122/ccs | No overlap | 2856 |
| m161203_032609_00127_c101108862550000001823246505011795_s1_p0/149838/ccs | 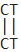 | 3022 |
| m161203_032609_00127_c101108862550000001823246505011795_s1_p0/146657/ccs | 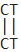 | 3022 |
| m161203_032609_00127_c101108862550000001823246505011795_s1_p0/149045/ccs | 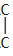 | 3479 |
| m161203_032609_00127_c101108862550000001823246505011795_s1_p0/125602/ccs | 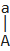 | 1298 |
| m161203_032609_00127_c101108862550000001823246505011795_s1_p0/125602/ccs | 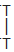 | 2827 |
| m161203_032609_00127_c101108862550000001823246505011795_s1_p0/42569/ccs | 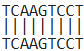 | 1178 |
| m161203_032609_00127_c101108862550000001823246505011795_s1_p0/42569/ccs | 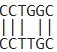 | 1902 |
| m161203_032609_00127_c101108862550000001823246505011795_s1_p0/137289/ccs | 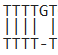 | 2631 |
| m161203_032609_00127_c101108862550000001823246505011795_s1_p0/122326/ccs | 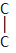 | 3623 |

**Table S2**

**PCR leaping products from previous CRISPR editing study 2. Genome editing with the HDR-enhancing DNA-PKcs inhibitor AZD7648 causes large-scale genomic alterations**

The nanopore amplicon sequencing results of the negative control used in Figure 1c (Figure1c_RPE1p53neg_GAPDH_Unedited, SRR31093017) were downloaded from the NCBI Sequence Read Archive. The FASTQ file was filtered by keeping only the reads that had both primers extended by at least 10 bp using custom Python scripts with the following commands:

py trimrightend.py SRR31093017.fastq ggagctgtcagatgagacat SRR31093017_right_trimmed.fastq -m 2

py trimleftend.py SRR31093017_right_trimmed.fastq aggacaggcaacttggcaaa SRR31093017_both_trimmed.fastq -m 2

and

py trimrightend.py SRR31093017.fastq TTTGCCAAGTTGCCTGTCCT SRR31093017_r_right_trimmed.fastq -m 2

py trimleftend.py SRR31093017_r_right_trimmed.fastq ATGTCTCATCTGACAGCTCC SRR31093017_r_both_trimmed.fastq -m 2

The results were then mapped to the reference sequences and visualized in IGV. From the 20,007 reads, we found the following 13 PCR leaping products:

| Read | Junction | Deletion size (bp) |
| --- | --- | --- |
| SRR31093017.1.12033 | 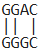 | 57 |
| SRR31093017.1.22307 | No overlap | 248 |
| SRR31093017.1.24851 | No overlap | 30 |
| SRR31093017.1.35439 | 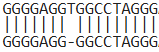 | 1875 |
| SRR31093017.1.14726 | No overlap | 4302 |
| SRR31093017.1.27602 | 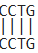 | 150 |
| SRR31093017.1.34337 | No overlap | 196 |
| SRR31093017.1.2513 | 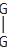 | 158 |
| SRR31093017.1.4075 | No overlap | 322 |
| SRR31093017.1.22462 | No overlap | 269 |
| SRR31093017.1.8702 | No overlap | 138 |
| SRR31093017.1.35431 | 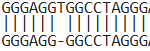 | 1876 |
| SRR31093017.1.26369 | 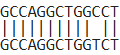 | 2913 |

**Table S3 PCR leaping products from previous CRISPR editing study 3. Single-cell multiplex approaches deeply map ON-target CRISPR-genotoxicity and reveal its mitigation by palbociclib and long-term engraftment**

The nanopore amplicon sequencing results of the negative control used in supplementary Figure 1 (SRR36268303) were downloaded from the NCBI Sequence Read Archive. The FASTQ file was filtered by keeping only the reads that had both primers extended by at least 10 bp using custom Python scripts with the following commands:

py trimrightend.py SRR36268303_1.fastq agaccagaaagatacatgtt SRR36268303_right_trimmed.fastq -m 2

py trimleftend.py SRR36268303_right_trimmed.fastq tgcagtagtcattcgtttat SRR36268303_both_trimmed.fastq -m 2

and

py trimrightend.py SRR36268303_1.fastq ATAAACGAATGACTACTGCA SRR36268303_r_right_trimmed.fastq -m 2

py trimleftend.py SRR36268303_r_right_trimmed.fastq AACATGTATCTTTCTGGTCT SRR36268303_r_both_trimmed.fastq -m 2

The results were then mapped to the reference sequence and visualized in IGV. From the 1159 reads, we found the following 27 PCR leaping products with short or no overlap at the junction (other 410 products have deletions around 4.6 kb. The 9.5 kb target area was found to contain two direct homologous repeats of 3.6 kb each. The large deletions can be explained by both the PCR leaping mechanism and the template switching mechanism. ):

| Read | Junction | Deletion size (bp) |
| --- | --- | --- |
| SRR36268303.11760 | 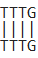 | 173 |
| @SRR36268303.9572 | 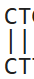 | 709 |
| SRR36268303.21623 | 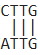 | 161 |
| SRR36268303.20235 | 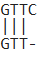 | 316 |
| SRR36268303.33973 | 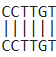 | 411 |
| SRR36268303.24238 | 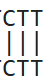 | 1327 |
| SRR36268303.27476 | 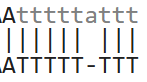 | 5841 |
| SRR36268303.35421 | 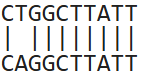 | 9004 |
| SRR36268303.12729 | 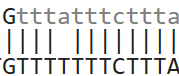 | 9194 |
| SRR36268303.25977 | 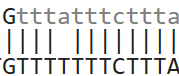 | 9194 |
| SRR36268303.14658 | 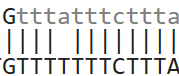 | 9194 |
| SRR36268303.12678 | 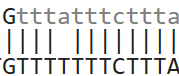 | 9194 |
| SRR36268303.8662 | 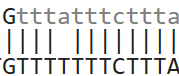 | 9194 |
| SRR36268303.15014 | 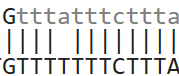 | 9194 |
| SRR36268303.37477 | 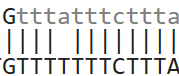 | 9194 |
| SRR36268303.18069 | 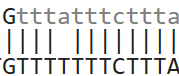 | 9194 |
| SRR36268303.25914 | 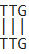 | 447 |
| SRR36268303.20634 | 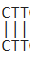 | 209 |
| SRR36268303.37435 | No overlap | 123 |
| SRR36268303.36785 | 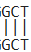 | 5137 |
| SRR36268303.9245 | 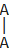 | 5125 |
| SRR36268303.20634 | 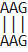 | 5207 |
| SRR36268303.16011 | 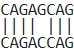 | 5041 |
| SRR36268303.4818 | 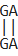 | 6485 |
| SRR36268303.24662 | 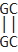 | 6178 |
| SRR36268303.16748 | 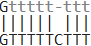 | 5907 |
| SRR36268303.7675 | 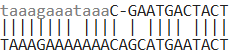 | 9189 |

**Table S4 PCR programs used in this study**

OnetaqTD40

| Initial Denaturation | 94°C | 5 min |
| --- | --- | --- |
| 28 cycles | 94°C 68°C to 40°C at -1 per cycle  68°C | 30 s  5 s  10 s |
| 40 Cycles | 94°C 55°C  68°C | 30 s  5 s  10 s |
| Final Extension | 68°C | 10 minutes |

OnetaqTD60

| Initial Denaturation | 94°C | 5 min |
| --- | --- | --- |
| 8 cycles | 94°C 68°C to 60°C at -1 per cycle  68°C | 30 s  15 s  30 s |
| 40 Cycles | 94°C 60°C  68°C | 30 s  15 s  30 s |
| Final Extension | 68°C | 10 minutes |

OnetaqTD50

| Initial Denaturation | 94°C | 5 min |
| --- | --- | --- |
| 18 cycles | 94°C 68°C to 50°C at -1 per cycle  68°C | 30 s  15 s  30 s |
| 40 Cycles | 94°C 50°C  68°C | 30 s  15 s  30 s |
| Final Extension | 68°C | 10 minutes |

OnetaqTD40a

| Initial Denaturation | 94°C | 5 min |
| --- | --- | --- |
| 28 cycles | 94°C 68°C to 40°C at -1 per cycle  68°C | 30 s  15 s  30 s |
| 40 Cycles | 94°C 50°C  68°C | 30 s  15 s  30 s |
| Final Extension | 68°C | 10 minutes |

Q5TD40

| Initial Denaturation | 98°C | 3 min |
| --- | --- | --- |
| 50 cycles | 98°C 65°C to 40°C at -0.5 per cycle  72°C | 10 s  15 s  10 s |
| 30 Cycles | 98°C 58°C  68°C | 20 s  15 s  10 s |
| Final Extension | 72°C | 10 minutes |

TaqTD50

|  | 95°C | 3 min |
| --- | --- | --- |
| 18 cycles | 95°C 68°C to 50°C at -1 per cycle  72°C | 30 s  10 s  5 s |
| 50 Cycles | 95°C 50°C  72°C | 30 s  10 s  5 s |
| Final Extension | 72°C | 10 minutes |

Taq50

|  | 95°C | 3 min |
| --- | --- | --- |
| 68 Cycles | 95°C 50°C  72°C | 30 s  10 s  5 s |
| Final Extension | 72°C | 10 minutes |

DreamtaqTD20

| Initial Denaturation | 95°C | 5 min |
| --- | --- | --- |
| 52 cycles | 95°C 62°C to 20°C at -0.8 per cycle  72°C | 30 s  50 s  60 s |
| 30 Cycles | 95°C 58°C  72°C | 30 s  30 s  60 s |
| Final Extension | 72°C | 10 minutes |

DreamtaqTD50

| Initial Denaturation | 95°C | 5 min |
| --- | --- | --- |
| 22 cycles | 95°C 72°C to 50°C at -1 per cycle  72°C | 30 s  10 s  10 s |
| 40 Cycles | 95°C 65°C  72°C | 30 s  10 s  10 s |
| Final Extension | 72°C | 10 minutes |

DreamtaqTD66

| Initial Denaturation | 95°C | 3 min |
| --- | --- | --- |
| 12 cycles | 95°C 72°C to 66°C at -0.5 per cycle  72°C | 30 s  30 s  9 min |
| 37 Cycles | 95°C 66°C  72°C | 30 s  30 s  9 min |
| Final Extension | 72°C | 10 minutes |

OnetaqTD40b

| Initial Denaturation | 94°C | 5 min |
| --- | --- | --- |
| 10 Cycles | 94°C 65°C  68°C | 30 s  5 s  10 s |
| 50 cycles | 94°C 65°C to 40°C at -0.5 per cycle  68°C | 30 s  50 s  10 s |
| 30 Cycles | 94°C 65°C  68°C | 30 s  5 s  20 s |
| Final Extension | 68°C | 10 minutes |

**Table S5 Primers used in this study and their sequences**

| primer | sequence |
| --- | --- |
| PUC1 | AATGGTTTCTTAGACGTCAGGTG |
| PUC2 | GCGGTTTGCGTATTGGG |
| PUC3 | GCATTTTGCCTTCCTGTTTT |
| PUC4 | GGGGATCCTCTAGAGTCGACC |
| PUC5 | TTCCTGTTTTTGCTCACCCA |
| PUC6 | GTTATCCGCTCACAATTCCAC |
| PUC7 | GGATGGCATGACAGTAAGAGAA |
| PUC8 | ATTTAGAAAAATAAACAAATAGGGGT |
| PUC9 | GGGAGTCAGGCAACTATGGAT |
| PUC10 | AATAAGGGCGACACGGAAAT |
| PUC11 | AAGCCATACCAAACGACGAG |
| PUC12 | GCAAAAACAGGAAGGCAAAAT |
| PUC13 | CCGCCTACATACCTCGCTCT |
| PUC14 | CAACTTTATCCGCCTCCATC |
| PUC15 | GCCACCACTTCAAGAACTCTGT |
| PUC16 | CTCCTTCGGTCCTCCGAT |
| PUC17 | GTTATCCCCTGATTCTGTGGAT |
| PUC18 | CGGAAAAAGAGTTGGTAGCTCT |
| PUC19 | TGAGCGAGGAAGCGGAAG |
| PUC20 | GAGCGAGGTATGTAGGCGG |
| lambda1 | ATGGCATTCTCTGGTTTTCGT |
| lambda2 | CCGCTGCTCCTGACTGTTC |
| lambda3 | GGAACTGCGACTGGATAGGC |
| lambda4 | TTTCTCCGTGGTGAAGGGATA |
| lambda5 | TTTCTGGGTAGATCGGGTTTC |
| lambda6 | TGTCTGTTCAGGGGCATTATT |
| lambda7 | GCTTTGGTGATGGCTATTCTC |
| lambda8 | TTACCGTTATCCGTTGTCTGTAT |
| lambda9 | GGAAACCACATACCGCTTCAC |
| lambda10 | GCCATATTCACCCCACAAAAA |
| lambda11 | CTGGACAATCAAGGGGAAAC |
| lambda12 | CAATACGATAGAAAAACAAGGCG |
| lambda13 | TTTACCCGTCCTTGGGTCC |
| lambda14 | TATTCTGTGTGCTTATGCTTGC |
| lambda15 | GAATGTAGTGGCGGAAAAGG |
| lambda16 | GAATGTGTAAGAGCGGGGTT |
| lambda17 | CAAGCAACAGGCAGGCGT |
| lambda18 | CAGTATTTGGTGAAGGGAACGA |
| lambda19 | GGTAGTGAGATGAAAAGAGGCG |
| lambda20 | CTGTGGACATAGTTAATCCGGG |
| Xj527.1 | CGTCGACATTTAGGTGACACTATAGAA |
| XJ528.1 | CGCTAATACGACTCACTATAGGGAGA |
| XJ462 | TGCAGTAGCGGATCACCGTTGCCCCTTGGCGC |
| xl29 | ctccaccgacacggtgtcgg |
| xl30 | GCGCTGTGCTCGCCGACGAC |
| Xj527.2 | TCTCCCGTCGACATTTAGGTGACACTATAGAA |
| XJ528.2 | TTCTATCGCTAATACGACTCACTATAGGGAGA |
| PUC13 | CCGCCTACATACCTCGCTCT |
| PUC14 | CAACTTTATCCGCCTCCATC |
| Xj527.7 | TTCTAatACTCGTCGACATTTAGGTGACACTATAGAA |
| XJ528.7 | TCTCaaAGTGCTAATACGACTCACTATAGGGAGA |
| xj595 | aaaaaaCCTGAAGTCAGCCCCATACG |
| xj596 | aaaaaaCAGCCTAATTACCCTGTTATCCC |
| 1s | CTTGCCCGCGAAGAAGCCCG |
| 1a | ACCGTGCACATCATGGCCCACC |
| 2s | TTACAGCACCTGGCGCAGGAATG |
| 2a | TTGCCGAGATCAACACCGACACC |
| 3s | GTCTTTCGGCGCAGCGAAGAATG |
| 3a | TCAACGATGATGACTGGACCTGGGAG |
| 4s | TGCCATGGTCGAAGAACAGCACG |
| 4a | CTCGCCGCCAACCCCGGTAT |
| 5s | CCATTTCATGGGTGACGCAGACC |
| 5a | TGGAAGACGCCTACGACAACGACC |
| 6s | CCAGCACCTCACCGACCATTTCC |
| 6a | GCCACCGTTACCGACAGAGCACG |
| 7s | CAGCATCACTTTCGGGTCCATCG |
| 7a | GGCCTGGTAGACCACATCTGCCC |
| 8s | CTGTTGCTGGCCGCCAGACAAGC |
| 8a | CGCGCTGTCGGTGTCAGAAAACG |
| 9s | TGCGAAATACCCACCTTCTCCAGC |
| 9a | AGGTTGCCCCATCCCTCAGCGT |
| 10s | GTTGCGCTTGCGCACCACCTTCT |
| 10a | AACACCCACAGACATCGGGACTGC |
| 1 | aaaaaaGCCGTCGACATTTAGGTGACACTATAGAA |
| 3 | aaaaggGCCGTCGACATTTAGGTGACACTATAGAA |
| 5 | aaaattGCCGTCGACATTTAGGTGACACTATAGAA |
| 7 | aaaaccGCCGTCGACATTTAGGTGACACTATAGAA |
| 9 | aaggaaGCCGTCGACATTTAGGTGACACTATAGAA |
| 11 | aaggttGCCGTCGACATTTAGGTGACACTATAGAA |
| 13 | aattaaGCCGTCGACATTTAGGTGACACTATAGAA |
| 15 | aattggGCCGTCGACATTTAGGTGACACTATAGAA |
| 17 | aattttGCCGTCGACATTTAGGTGACACTATAGAA |
| 19 | aattccGCCGTCGACATTTAGGTGACACTATAGAA |
| 21 | aaccaaGCCGTCGACATTTAGGTGACACTATAGAA |
| 23 | aaccttGCCGTCGACATTTAGGTGACACTATAGAA |
| 2 | ttaaaaGCCGTCGACATTTAGGTGACACTATAGAA |
| 4 | ttaaggGCCGTCGACATTTAGGTGACACTATAGAA |
| 6 | ttaattGCCGTCGACATTTAGGTGACACTATAGAA |
| 8 | ttaaccGCCGTCGACATTTAGGTGACACTATAGAA |
| 10 | ttggaaGCCGTCGACATTTAGGTGACACTATAGAA |
| 12 | ttggttGCCGTCGACATTTAGGTGACACTATAGAA |
| 14 | ttttaaGCCGTCGACATTTAGGTGACACTATAGAA |
| 16 | ttttggGCCGTCGACATTTAGGTGACACTATAGAA |
| 18 | ttttttGCCGTCGACATTTAGGTGACACTATAGAA |
| 20 | ttttccGCCGTCGACATTTAGGTGACACTATAGAA |
| 22 | ttccaaGCCGTCGACATTTAGGTGACACTATAGAA |
| 24 | ttccttGCCGTCGACATTTAGGTGACACTATAGAA |
| A | aaaaaaGGCCGCTAATACGACTCACTATAGGGAGA |
| C | aaaaggGGCCGCTAATACGACTCACTATAGGGAGA |
| E | aaaattGGCCGCTAATACGACTCACTATAGGGAGA |
| G | aaaaccGGCCGCTAATACGACTCACTATAGGGAGA |
| I | aaggaaGGCCGCTAATACGACTCACTATAGGGAGA |
| K | aaggttGGCCGCTAATACGACTCACTATAGGGAGA |
| M | aattaaGGCCGCTAATACGACTCACTATAGGGAGA |
| O | aattggGGCCGCTAATACGACTCACTATAGGGAGA |
| B | aattttGGCCGCTAATACGACTCACTATAGGGAGA |
| D | aattccGGCCGCTAATACGACTCACTATAGGGAGA |
| F | aaccaaGGCCGCTAATACGACTCACTATAGGGAGA |
| H | aaccttGGCCGCTAATACGACTCACTATAGGGAGA |
| J | ttaaaaGGCCGCTAATACGACTCACTATAGGGAGA |
| L | ttaaggGGCCGCTAATACGACTCACTATAGGGAGA |
| N | ttaattGGCCGCTAATACGACTCACTATAGGGAGA |
| P | ttaaccGGCCGCTAATACGACTCACTATAGGGAGA |
